# Common non-antibiotic drugs enhance selection for antimicrobial resistance in mixture with ciprofloxacin

**DOI:** 10.1101/2025.03.23.644574

**Authors:** April Hayes, Lihong Zhang, Jason Snape, Ed Feil, Barbara Kasprzyk-Hordern, William H Gaze, Aimee K Murray

## Abstract

Antimicrobial resistance (AMR) is a major health concern, and a range of antibiotic and non-antibiotic agents can select for AMR across a range of concentrations. Selection for AMR is often investigated using single compounds, however, in the natural environment and the human body, pharmaceuticals will be present as mixtures, including both non-antibiotic drugs (NADs), and antibiotics. Here, we assessed the effects of one of three NADs in combination with ciprofloxacin, a commonly used antibiotic that is often found at concentrations in global freshwaters sufficiently high to select for AMR. We used a combination of growth assays and qPCR to determine selective concentrations of mixtures and used metagenome sequencing to identify changes to the resistome and community composition. The selective concentration of ciprofloxacin was reduced from 40µg/L to 10µg/L in the mixtures, and mixtures showed a stronger selection for some AMR genes such as *tolC* and *qnrB*. The communities exposed to the mixtures also showed changed community compositions. These results demonstrate that NADs and ciprofloxacin are more selective than ciprofloxacin alone, and these mixtures can cause distinct changes to the community composition. This indicates that future work should consider combinations of antibiotics and NADs as drivers of AMR when considering its maintenance and acquisition.

## Introduction

Antimicrobial resistance (AMR) is a global health threat, with 1.27 million deaths in 2019 directly caused by antibiotic resistant bacterial infections [1]. Traditionally, resistance to antibiotics has been determined by identifying concentrations that inhibit growth. However, research shows that low concentrations of antibiotics can select for antibiotic resistance in both single species [2, 3], and in bacterial communities [4–9]. Additionally, other non-antibiotic compounds can co-select for antibiotic resistance, including metals and biocides [10, 11]. Non-antibiotic drugs (NADs) have previously been shown to reduce bacterial growth [12, 13], increase horizontal gene transfer rates [14–17], and select for antibiotic resistance [18–20], in single species experiments at therapeutic concentrations. There is some evidence suggesting that NADs at lower, more environmentally relevant concentrations may not select, or select less strongly for AMR, in both single species and mixed communities [21, 22]. We previously tested three NADs commonly found in the environment - diclofenac, metformin and 17-β-estradiol – on their selective potential for AMR [22]. Diclofenac, a non-steroidal anti-inflammatory drug [32], is one of the five most frequently detected pharmaceuticals in the aquatic environment [23]. Metformin is used in front-line diabetes treatments [24] and 17-β-estradiol is a naturally produced hormone but is also used in hormone replacement therapy [25, 26]. Diclofenac, metformin and 17-β-estradiol did not strongly select for antibiotic resistance genes within a bacterial community, or affect bacterial diversity [22]. However, they did have antimicrobial activity in terms of impact on growth rate and additionally had some effects on metal resistance gene abundance/diversity [22].

Pharmaceuticals are present in both the human body and in the aquatic environment at a range of concentrations [24, 27–30]. NADs will be present alongside antibiotics in these environments as both simple and complex mixtures of multiple pharmaceuticals [31]. Environmentally relevant concentrations of ciprofloxacin can select for AMR [2, 9] and previous environmental risk assessments have shown that there is a risk of AMR selection by ciprofloxacin in various wastewater environments, even in high income countries [32].

Several studies have studied the effects of mixtures of NADs and antibiotics with a specific focus on their capacity to reduce or completely inhibit bacterial growth or increase antibiotic susceptibility [33, 34]. For example, diclofenac and metformin have been shown to act both synergistically and antagonistically with a range of antibiotics across a range of classes and species [35] including the pathogen *Pseudomonas aeruginosa*. Diclofenac can also increase the inhibitory activity of ciprofloxacin in *Proteus mirabilis* [36]. Alternatively, metformin has been shown to increase the inhibitory activity of tetracycline antibiotics, and restored tetracycline susceptibility in a resistant *Escherichia coli* strain [37]. Overall, there is not enough evidence to conclusively suggest mechanistic insights into combinations of these pharmaceuticals. These results may be species specific, or related to altered gene expression, and there is less evidence when considering their effects on complex communities.

Furthermore, most research has focused on testing the effects of mixtures of antibiotics. Work in this area has illustrated within-species variability in responses to antibiotic mixtures [38], and shown that the responses of single species do not predict the responses of a more complex community [39]. Furthermore, there is evidence to suggest that complex microbial communities may be more resilient to mixtures than individual species [39], which also occurs with single antibiotic compounds [40]. Additionally, research has shown that when antimicrobials are combined in complex mixtures, a variety of interactions can occur. This includes increased and decreased evolution of resistance [41–43]. Overall, mixtures are likely to lead to different selection dynamics in bacterial communities compared to single compounds alone due to increased variability in both species and genes.

In this study, we aimed to understand the effects of simple mixtures in a complex microbial community, since this is first step to understanding complex mixture effects. We experimentally spiked single concentrations of diclofenac, metformin, and 17-β-estradiol alongside a range of ciprofloxacin concentrations to see if this affected the minimal selective concentration of ciprofloxacin. Firstly, we determined if there was a significant reduction in the growth of the community in the mixture compared to ciprofloxacin alone, and secondly, if the minimal selective concentration of ciprofloxacin changed in the presence of the NAD. We used *intI1* as the selective concentration endpoint, since this has previously been shown to increase with ciprofloxacin selection [9], and *intI1* has been suggested as a proxy for antimicrobial resistance acquisition in environmental monitoring [44]. Finally, we determined whether the community resistome or composition changed in the mixtures using metagenome sequencing.

## Methods and Materials

### Pharmaceuticals

Diclofenac (Sigma Aldritch), metformin (Enzo), 17-β-estradiol (Sigma Aldritch), and ciprofloxacin (Sigma Aldritch) were acquired and dissolved in water, water, ethanol, and 0.8mol HCl and 1.2mL water respectively and filter sterilised. Aliquots were kept at −20°C for up to two weeks before use. Stock concentrations of pharmaceuticals were diluted in filter sterilised water to concentrations for use.

### Wastewater influent

Wastewater influent was collected from Falmouth (UK) wastewater treatment plant, in June 2022. The wastewater was collected in clean 1L glass Duran bottles and processed the same day. Wastewater was mixed 1:1 with 40% glycerol and kept frozen at −70°C until use. A wastewater bacterial community was used since it contains a large number of bacterial species, including those that are associated with humans, and can contain opportunistic pathogens [22]. Furthermore, this type of community is likely exposed to antimicrobials within the environment as highlighted earlier, so provides real-world relevance to this study.

### Growth Assays

Combinations of ciprofloxacin and either diclofenac, metformin, or 17-β-estradiol were tested across a range of ciprofloxacin concentrations with a spiked concentration of the NAD (diclofenac: 50µg/L and 25µg/L, metformin: 26µg/L and 13µg/L, 17-β-estradiol: 24.8µg/L and 12.4µg/L). The higher set of these concentrations had previously been shown to significantly reduce growth compared to a no-NAD control, and the lower concentration did not significantly reduce growth compared to a no-NAD control, within the same experimental set up [45]. All mixtures contained NAD at all ciprofloxacin concentrations including 0 µg/L.

A 96 well plate was filled with 180µL Iso-Sensitest broth (Oxoid). For the ciprofloxacin gradient, 180µL Iso-Sensitest broth with 15.6µg/L ciprofloxacin was added to the top 12 wells of the plate, and 180µL was serially diluted down the plate, leaving one row as a no-antibiotic control. The higher NAD concentration was added to four columns, the lower NAD concentration to four columns, and the remaining four columns were left as a no-NAD control (Figure 1). Wastewater influent was thawed, and washed with 0.85% NaCl twice to remove contaminants and nutrient carry over, and 10µL of this was inoculated into each well. The plate was sealed with a MicroAmp Optical seal and optical density was measured at 600nm (OD600) every ten minutes for 24 hours in a BioTek plate reader (Agilent), with five seconds of shaking at 180 rpm every ten minutes.

**Figure 1.**
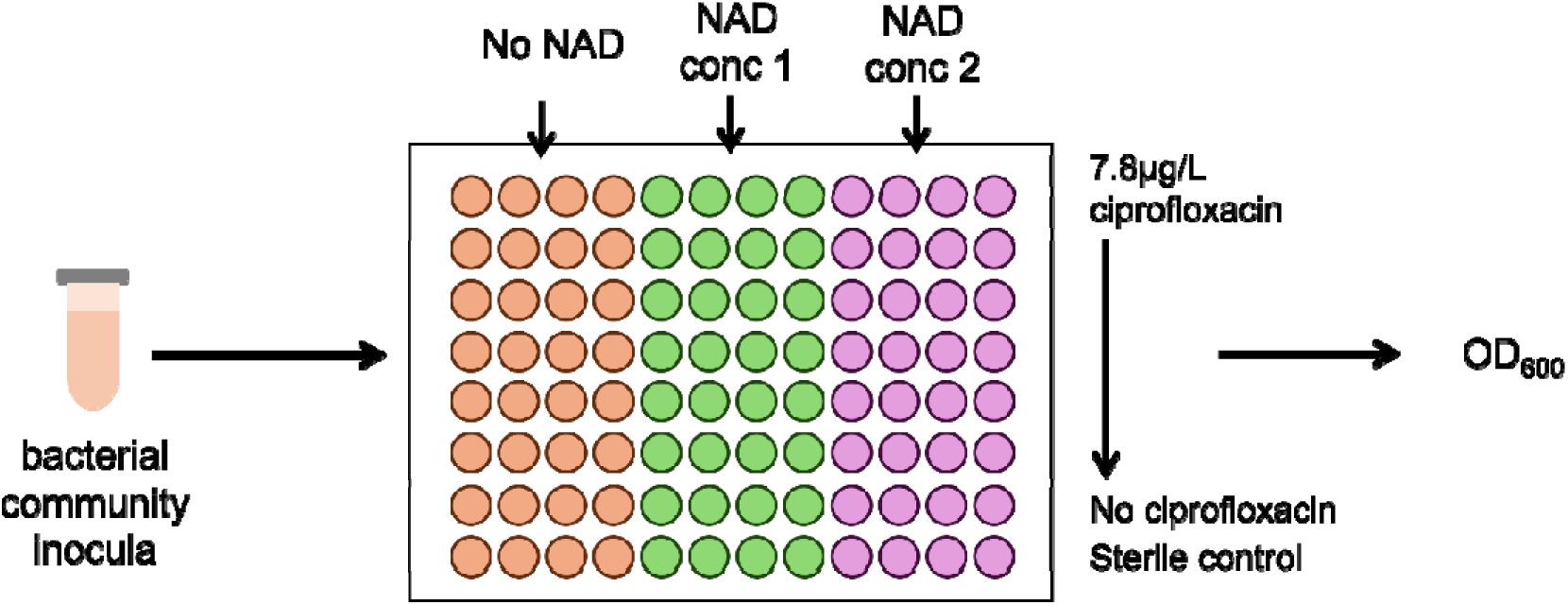
96 well plate layout for the growth assays for mixture experiments.

### Selection Experiments for qPCR analysis

Selection experiments were carried out across a ciprofloxacin gradient, with the addition of single spiked NAD concentrations. Firstly, the lowest observed effect concentration (LOEC) of ciprofloxacin was determined. Ciprofloxacin was tested at 40µg/L, 20µg/L, 10µg/L, 5µg/L, and 2.5µg/L and 0 µg/L. This range was informed by a previous study where the LOEC was calculated to be 15.6µg/L [9]. Then, mixture experiments using the NADs were performed over this ciprofloxacin concentration range, but including either diclofenac spiked at 50µg/L, metformin at 26µg/L or 17-β-estradiol at 24.8µg/L [22]. All mixtures contained NAD at all ciprofloxacin concentrations, including 0 µg/L.

To set up the experiments, wastewater influent was thawed and washed twice with 0.85% NaCl and inoculated with 10% vol/vol into Iso-Sensitest broth. This inoculated broth was separated into 30mL aliquots, which were spiked with the ciprofloxacin and NAD concentrations. These 30mL aliquots were then separated into 5mL microcosms. Day zero samples were taken at this time point. For this, 1mL of each microcosm was taken, centrifuged at 2100rpm for two minutes and the pellet resuspended with 20% glycerol. Samples were kept at −70°C until use. The microcosms were incubated at 37°C with 180rpm shaking, with transfers into fresh media with fresh NAD and ciprofloxacin daily for seven days. On day seven, samples were taken by mixing 0.5mL culture 1:1 with 40% glycerol and kept at −70°C until use.

### QPCR analysis

QPCR was performed using day zero and day seven samples to determine effects on selection after the weeklong experiment. DNA was extracted using a DNeasy UltraClean Microbial Kit (Qiagen). All steps were carried out to manufacturer’s instructions with the initial centrifugation step extended to one minute. Extracted DNA was diluted 5X with TE and stored at 4°C before use. QPCR was performed using *intI1* primers and standardised to *16S* rRNA copy number using a QuantStudio 7 Real-Time PCR machine (Thermo Fisher). Prevalence was calculated by dividing the *intI1* copy number by the 16S rRNA copy number. The reaction mix included 10µL SYBR MasterMix with ROX and SYBR (PrimerDesign), 1µL forward primer (Integrated DNA Technologies), 1µL reverse primer (Integrated DNA Technologies), 0.2µL Bovine Serum Albumin and 2.8µL nuclease free water (Ambion). The cycling protocol was as follows – 120 second hold at 95°C, 50 rounds of cycling with 10 seconds at 95°C for denaturation and 60 seconds at 60°C for data collection. Only runs with an efficiency of 90-11%, and an R^2^ of greater than 0.99 were used in analyses. Primer and gblock sequences are presented in Supplementary Table 1.

### Metagenome Sequencing

For Illumina metagenome sequencing, day seven samples were thawed, and DNA extracted using a DNeasy UltraClean Microbial Kit, with all steps carried out to the manufacturer’s instructions save for two exceptions. The initial centrifugation step was elongated to 2 minutes, and the centrifugation of the PowerBead tubes was increased to one minute at 12,000G. Extracted DNA was purified using a standard RNase A and standard Ampure XP bead protocol. DNA was eluted in 10mM Tris-HCl and stored at 4°C until being sent for sequencing. NEB PCR-free library prep was carried out by the Exeter Sequencing Centre prior to sequencing using a NovaSeq SP to a depth of up to 20GB per sample.

### Metagenome Analyses

Trimmed reads from the Exeter Sequencing Service were used in all analyses. All reads were checked for quality using FastQC and MultiQC [46]. AMR++ was used to process the reads [47]. Low quality reads and reads mapping to host (human) were removed. For the resistome analysis, reads were aligned to the MEGARES 3.0 database [47], which includes multiple resistome databases including BacMet [48], ResFinder [49], and CARD [50]. For the microbiome analysis, *kraken2* [51] was used to identify taxonomy as part of the AMR++ pipeline using the minikraken database [52, 53]. Outputs from these pipelines were input into R, and converted into phyloseq objects using *phyloseq* v1.48.0 [54]. Reads were normalised using *metagenomeSeq* v1.46.0 with cumulative sum scaling [55, 56]. Relative abundances (proportions) were then created using these normalised data.

### Data Analysis

All statistical analyses were carried out in R version 4.4.1 [57]. All figures were generated using *ggplot2* 3.5.1 [58] and *MetBrewer* v0.2.0 [59]. For all models, the most parsimonious model was used, determined by sequentially deleting terms and comparing model fits using *X^2^* tests. Only those models with residuals fitting assumptions were used. Fit of residuals were checked using *DHARMa* v0.4.6 [60].

#### Growth Analyses

To determine the minimal selective concentrations or LOECs of the mixtures using growth, a method previously used was applied [61]. In summary, the time point in exponential phase with the largest dose response was determined using either Spearman’s or Pearson’s correlation test, as determined by the normality fit of the data. Then, at this time point, a Dunn’s test (*dunn.test* v1.3.5) was used to identify which concentrations significantly differed from the control growth (i.e. no ciprofloxacin and no NAD). The lowest concentration that was significantly different to the control was determined as the LOEC

To determine total growth capacity or productivity, total area under the curve (AUC) was used. AUC was determined using the *growthcurver* v0.3.1 package [62]. The AUC from exponential growth phase was used in linear mixed effect models using *lme4* v1.1.31 [63], with concentration of ciprofloxacin and treatment of NAD as fixed effects, and microcosm as a random effect. Pairwise comparisons were determined using *emmeans* v1.8.2 [64], and p values adjusted for multiple comparisons using false discovery rate.

#### QPCR Analyses

*IntI1* prevalence was calculated by dividing the *intI1* quantity by the *16S* rRNA quantity. These prevalences were used to calculate LOECs. LOECs were determined using linear mixed effect models, with time and treatment as fixed effects, and microcosm was included as a random effect. Pairwise comparisons were calculated as above. LOECs were determined to be the lowest concentration that showed a significantly increased *intI1* prevalence compared to the day seven control.

#### Diversity analyses

Alpha diversity or richness was used to identify the total number of taxa and genes present in each sample. *Phyloseq* was used to estimate Shannon’s index [65], which was used to identify the evenness of taxa and genes. Tests for significant differences in treatments for alpha diversity were tested using linear models, with richness as the response variable, and mixture type and ciprofloxacin concentration as explanatory variables. Pairwise comparisons were calculated as above. For beta diversity analyses, Bray-Curtis ordinations were calculated using *vegan* v2.6.6.1 [66] for both the resistome and the taxonomy. Changes to the diversity were determined using Analysis of Similarity (ANOSIM) tests.

#### Changes to resistance gene classes or species

Kruskal-Wallis tests were used to identify differences in gene classes, or taxonomic order by mixture type, using the normalised gene counts. Genes or taxonomic orders that differed significantly in at least one treatment were tested to determine if mixture type and/or ciprofloxacin concentration significantly affected gene abundance by linear models and ANOVAs.

#### Log2-fold change in resistance genes or species

For both the resistome and the community taxonomy, significant log2-fold changes between mixtures and the ciprofloxacin alone treatments were identified using *DESeq2* v 1.44.0 [67].

## Results

### NAD and ciprofloxacin mixtures significantly reduced community productivity

To test differences in community productivity, we tested changes to the AUC, which can be considered a proxy for overall productivity or growth capacity. The AUC of the mixtures were compared to in-plate AUC data of ciprofloxacin alone. Firstly, none of the lower concentrations of the NADs significantly reduced growth compared to the no-NAD control (p<0.05). However, all three NADs at the higher concentration in mixture with ciprofloxacin significantly reduced the productivity of communities compared to growth in the no NAD control (Figure 2), across the entire ciprofloxacin concentration gradient (p<0.001). We also found that as expected, total AUC reduced with increased ciprofloxacin concentration (p<0.001). More details of the outputs for these models are detailed in the Supplementary File, section 1.

**Figure 2.**
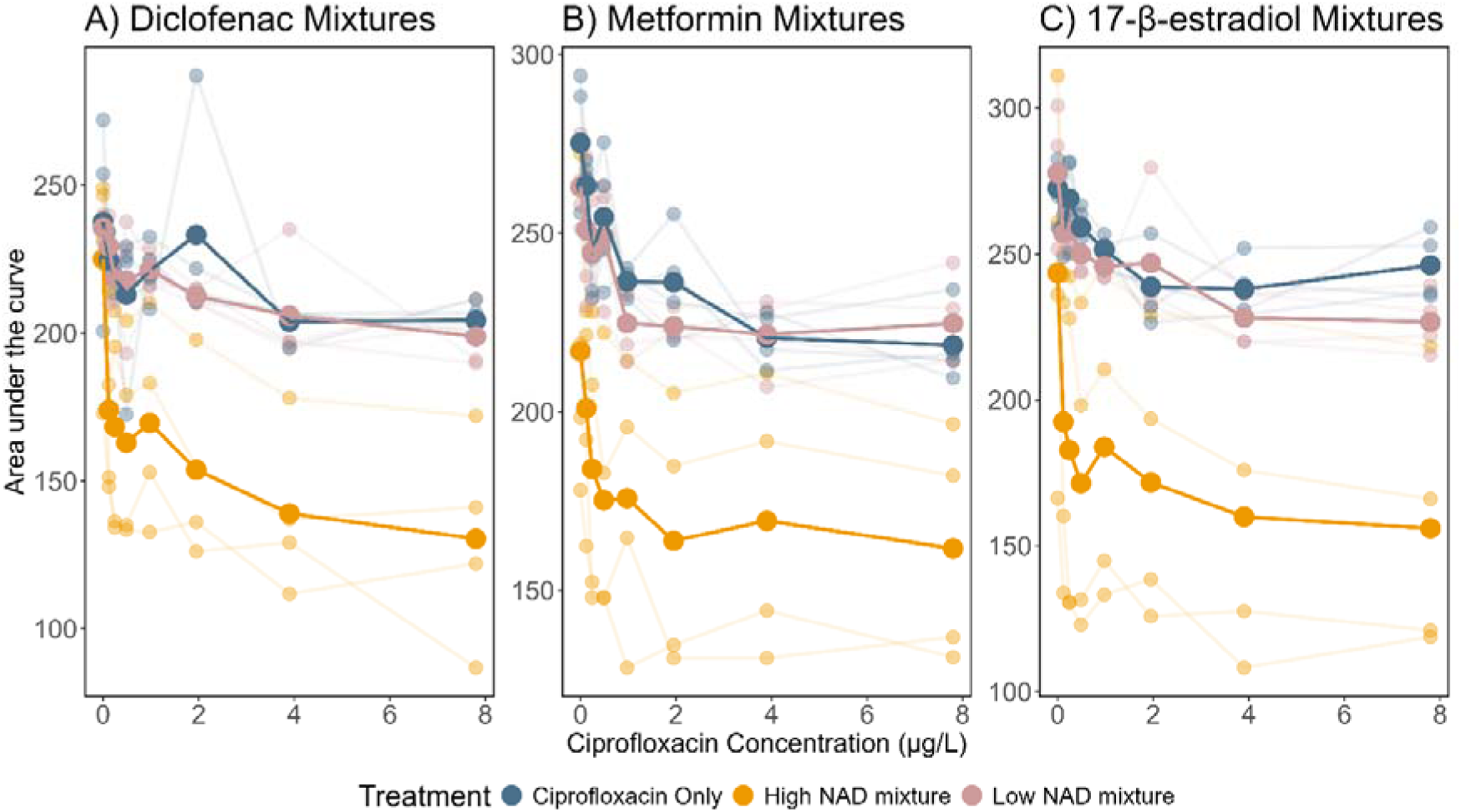
Community productivity (area under the curve) during exponential phase across a ciprofloxacin concentration gradient in mixture with A) diclofenac, B) metformin, and C) 17-β-estradiol. All low and high NAD mixtures include NAD at all ciprofloxacin concentrations, including 0ug/L ciprofloxacin. Pale linked points indicate individual replicates. Larger brighter points that are linked indicate the mean at each concentration.

Secondly, we determined the growth-based LOEC for the mixtures containing the higher NAD concentrations, using a low-cost method previously published [61]. Previous work has indicated that reduction in growth is the strongest indicator of selection for AMR [68]. We found that all mixtures reduced the estimated selective concentration of ciprofloxacin. The diclofenac-ciprofloxacin mixture reduced the LOEC of ciprofloxacin from 3.7µg/L to 0.12µg/L, a 32-fold decrease. The metformin-ciprofloxacin mixture reduced the LOEC of ciprofloxacin from 0.98µg/L to 0.24µg/L, a 4-fold decrease. The 17-β-estradiol-ciprofloxacin mixture reduced the LOEC of ciprofloxacin from 1.95µg/L to 0.24µg/L, an 8-fold decrease.

### NAD mixtures altered selection for *intI1* across ciprofloxacin concentrations

Next, we determined whether there was specific selection for the commonly used AMR marker *intI1.* We hypothesised that the mixtures of NADs and ciprofloxacin would have increased selectivity compared to ciprofloxacin alone.

Ciprofloxacin alone selected for *intI1* at 40µg/L (Figure 3A). We found that *intI1* prevalence significantly increased with treatment (treatment main effect, X^2^=18.35, df=4, p=0.0011) and that only 40µg/L ciprofloxacin had a day seven prevalence that was significantly greater than the prevalence at day seven control population (p=0.012).

**Figure 3.**
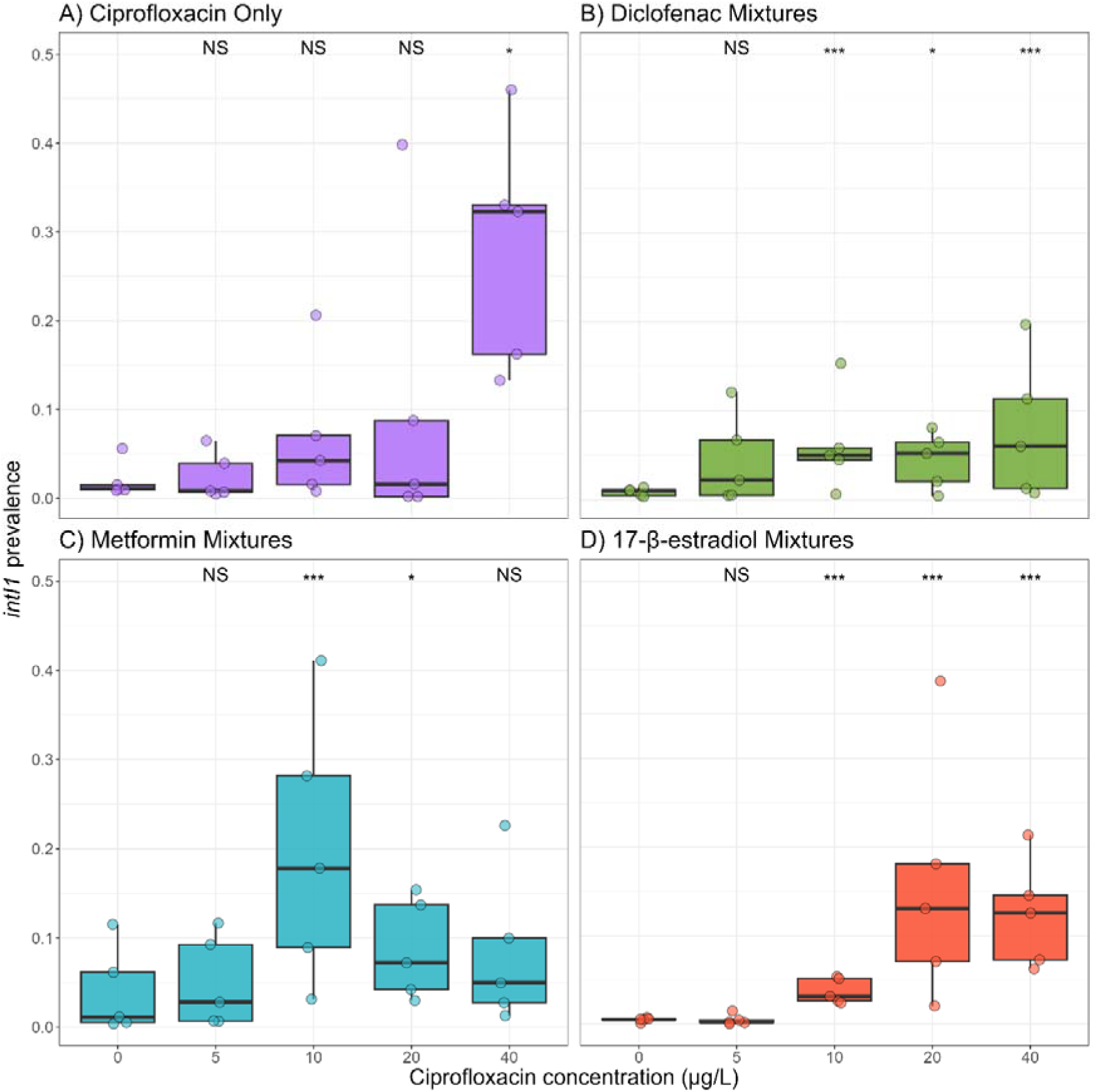
*IntI1* prevalence as a function of ciprofloxacin concentration in both ciprofloxacin alone (A) or in combination with diclofenac (B), metformin (C), or 17-β-estradiol (D). Five biological replicates shown. Significant differences of day seven prevalences are shown from pairwise comparisons to the day seven prevalence at 0µg/L. NS = non-significant, * = p<0.05, ** = p<0.01, *** = p<0.001.

We then tested the mixtures of NADs and ciprofloxacin and found that they all reduced the selective concentration to 10µg/L (Figure 3). In all cases, the concentration of *intI1* either increased or stayed constant with time (Supplementary Figure 1), so we only show day seven data in the following plots.

Firstly, 50µg/L diclofenac in mixture with ciprofloxacin (Figure 3B) reduced the selective concentration for *intI1*, with the communities exposed to 10µg/L (p=0.0085), 20µg/L (p=0.028) and 40µg/L (p=0.0085) showing significantly increased *intI1* prevalence compared to the control. Secondly, metformin at 26µg/L reduced the selective concentration of ciprofloxacin to 10µg/L (Figure 3C). We see that 10µg/L (p=0.0016), and 20µg/L (p=0.024) in the mixture significantly increased *intI1* prevalence, however this increase was not seen with 40µg/L ciprofloxacin (p=0.063). Finally, we found that 24.4µg/L 17-β-estradiol-ciprofloxacin mixture also reduced the selective concentration to 10µg/L (p<0.0001), with communities exposed to 20µg/L (p<0.0001) and 40µg/L (p<0.0001) also showing significantly increased *intI1* prevalence compared to the ciprofloxacin only community.

Despite mixtures of NADs and ciprofloxacin selecting for *intI1* at a lower ciprofloxacin concentration, the overall strength of selection in the mixtures was lower than that with ciprofloxacin alone, indicating some suppression of selection within the mixtures.

### Mixtures did not strongly alter total microbiome or resistome diversity

Next, we analysed the metagenomes of the evolved communities to understand how the mixtures might have affected the community composition and the resistome. We hypothesised that there would likely be increases in gene abundances occurring at lower ciprofloxacin concentrations in the mixtures, since the mixtures appeared to be more selective. We also hypothesised that it was likely that the mixtures had selected for different species, and that the mixtures would have decreased richness and diversity, since the communities were exposed to multiple pharmaceutical stressors.

There were no significant differences in richness of the taxa present in each sample. However, there were differences in the richness of the AMR genes between the treatments (Figure 4). The 17-β-estradiol mixture had a non-significant decrease in richness compared to the ciprofloxacin only treatment (p=0.071). Exposure to the metformin mixture significantly reduced resistome richness compared to the ciprofloxacin alone treatment (p=0.021). The diclofenac mixture showed a similar level of richness of AMR genes as the ciprofloxacin alone treatment.

**Figure 4.**
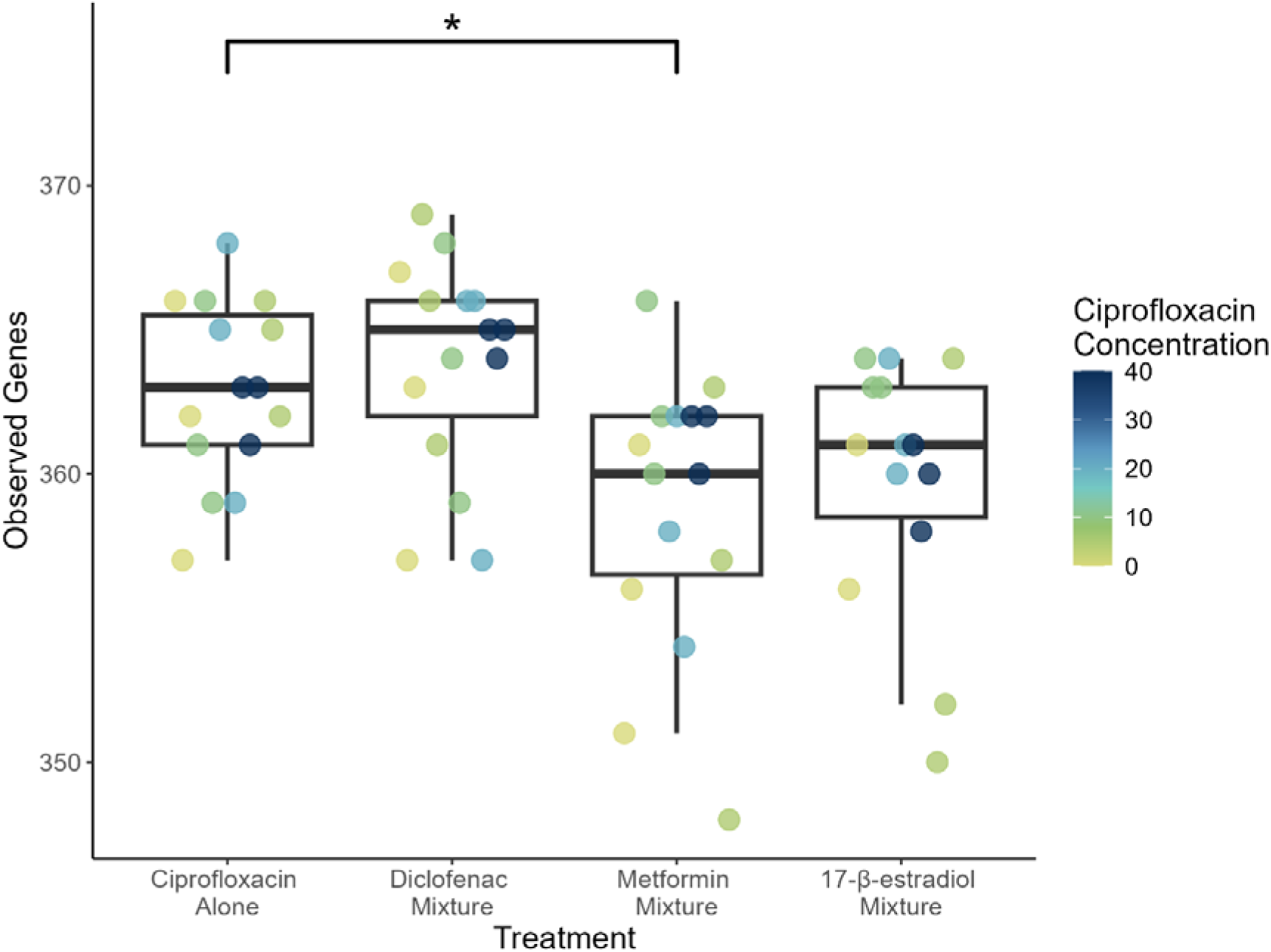
Resistome richness (number of observed AMR genes) of the evolved samples. Colour of the points indicates ciprofloxacin concentration. Asterisks indicate significance values where * = p<0.05, and ** = p<0.001.

There was no significant difference in the evenness of the resistome or the taxonomy between the mixture treatments. There was also no significant difference in the beta diversity of either the resistome or community composition (ANOSIM, p>0.05). (Supplementary Figures 2 and 3).

### Mixtures significantly altered the abundance of three AMR genes

We tested whether all fluroquinolone genes within the *AMR++* database had increased with treatment. We found that only *qnrB* had significantly altered abundance across all samples within each treatment (Supplementary Table 2). Next, we tested all genes within the resistome to see if they differed in at least one treatment and found that ten genes were significantly different in at least one sample (p<0.05) (Supplementary Table 3). Of these genes, six were significantly affected by mixture type either as main or interaction effect: *aph3-DPRIME, aph6, fecE, qnrB, tetA, tetQ,* and *tolC* (Figure 5). All details on the model outputs can be found in the Supplementary File in section 2.

**Figure 5.**
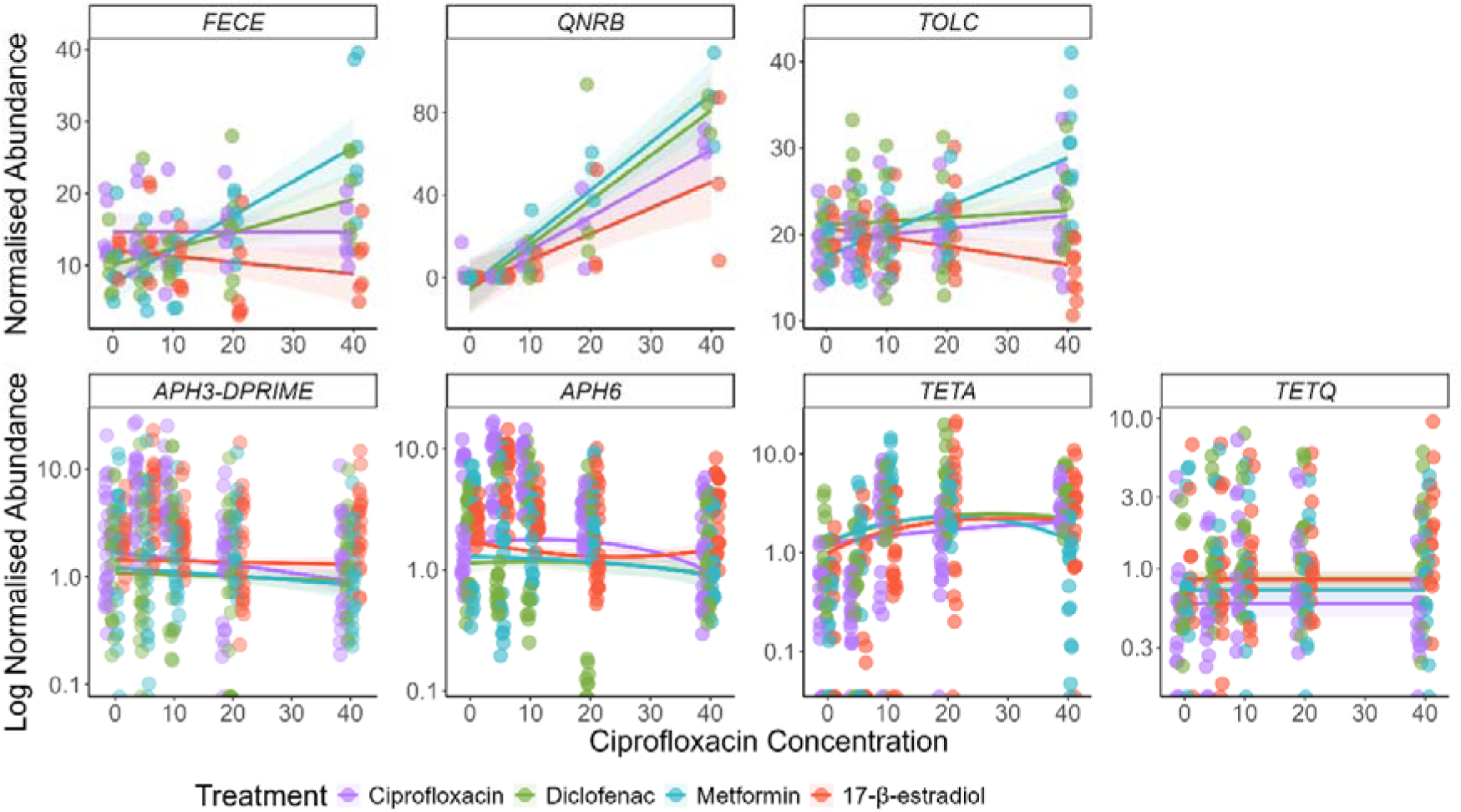
Normalised abundance of all genes that were significantly different in at least one sample and were significantly affected by NAD presence in the mixture. Colours indicate mixture treatments. Lines show best model fit, and the shaded areas show 95% confidence intervals. Points indicate the individual replicates.

*FecE*, *qnrB*, and *tolC* showed the largest changes in gene abundances. For these three genes, the same trends appeared with the metformin mixture showing the largest increase in gene abundances, followed by the diclofenac mixture. The 17-β-estradiol mixture either decreased in gene abundance (*fecE* and *tolC*) or showed the smallest increase (*qnrB*). For the other four genes, abundances changed very slightly and often had a unimodal response (e.g. *tetA*).

### Mixtures significantly altered the abundances of multiple bacterial taxa

We tested whether any order of microorganism was significantly different in at least one treatment. Two orders showed strong alterations across either concentration or mixture type: Caulobacterales and Nitrososphaerales. Caulobacterales abundance in the community decreased after exposure to ciprofloxacin only but increased in abundance in all the mixture treatments (interaction effect: F_3,110_=3.71, p=0.014) (Figure 6). Nitrososphaerales increased in abundance with ciprofloxacin concentration (concentration main effect: F_1,54_=17.52, p=0.00011), and had a decreased abundance in the mixtures compared to ciprofloxacin alone treatment (mixture main effect: F_3,54_=5.81, p=0.0016) (Figure 6).

**Figure 6.**
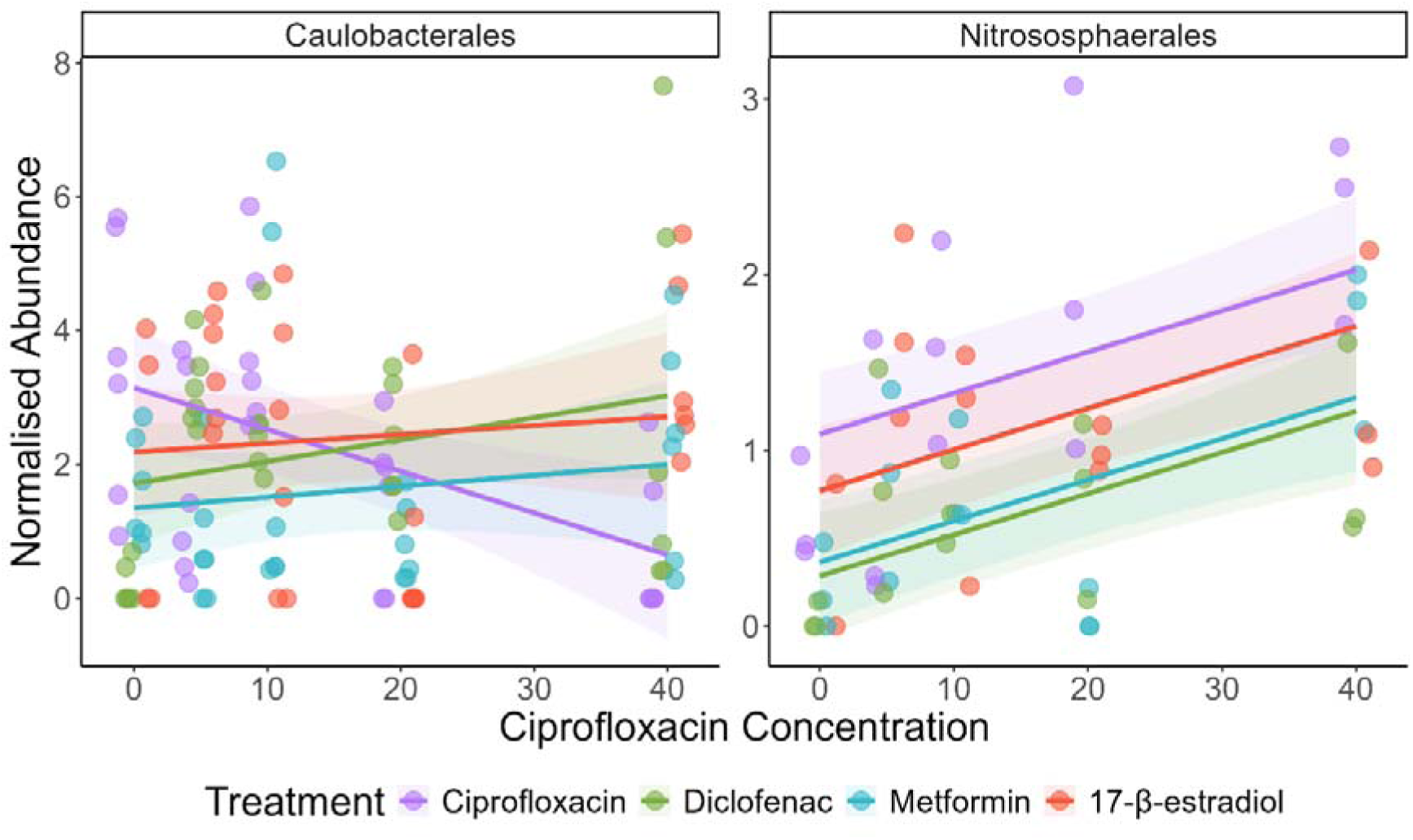
Normalised abundance of Caulobacterales and Nitrososphaerales as a function of ciprofloxacin concentration. Colours indicate mixture treatments. Lines show the best model fit, and shaded areas show 95% confidence intervals. Points represent the individual replicates.

Additionally, we grouped the community based on abundances greater than 10%. In these high abundance genera, there were no clear trends in changes to these genera, or phyla across the mixtures (Supplementary Figure 4). This indicates that was no dominance effect, and the communities were mostly stable across treatments.

However, over 100 species of microorganism had a log2-fold increase or decrease in each of the mixtures compared to the ciprofloxacin alone (Supplementary Figure 5). There were 43 species that either significantly increased or decreased in all three of the mixtures compared to the control (Figure 7). For these 43 species, the diclofenac and metformin mixtures showed the opposite response to the 17-β-estradiol mixture, i.e. if the diclofenac and metformin mixture showed an increased abundance of a particular species, the 17-β-estradiol mixture showed a decrease, and vice versa. Some of these species are known pathogens (e.g. *Pseudomonas. aeruginosa,* which increased in the 17-β-estradiol mixture).

**Figure 7.**
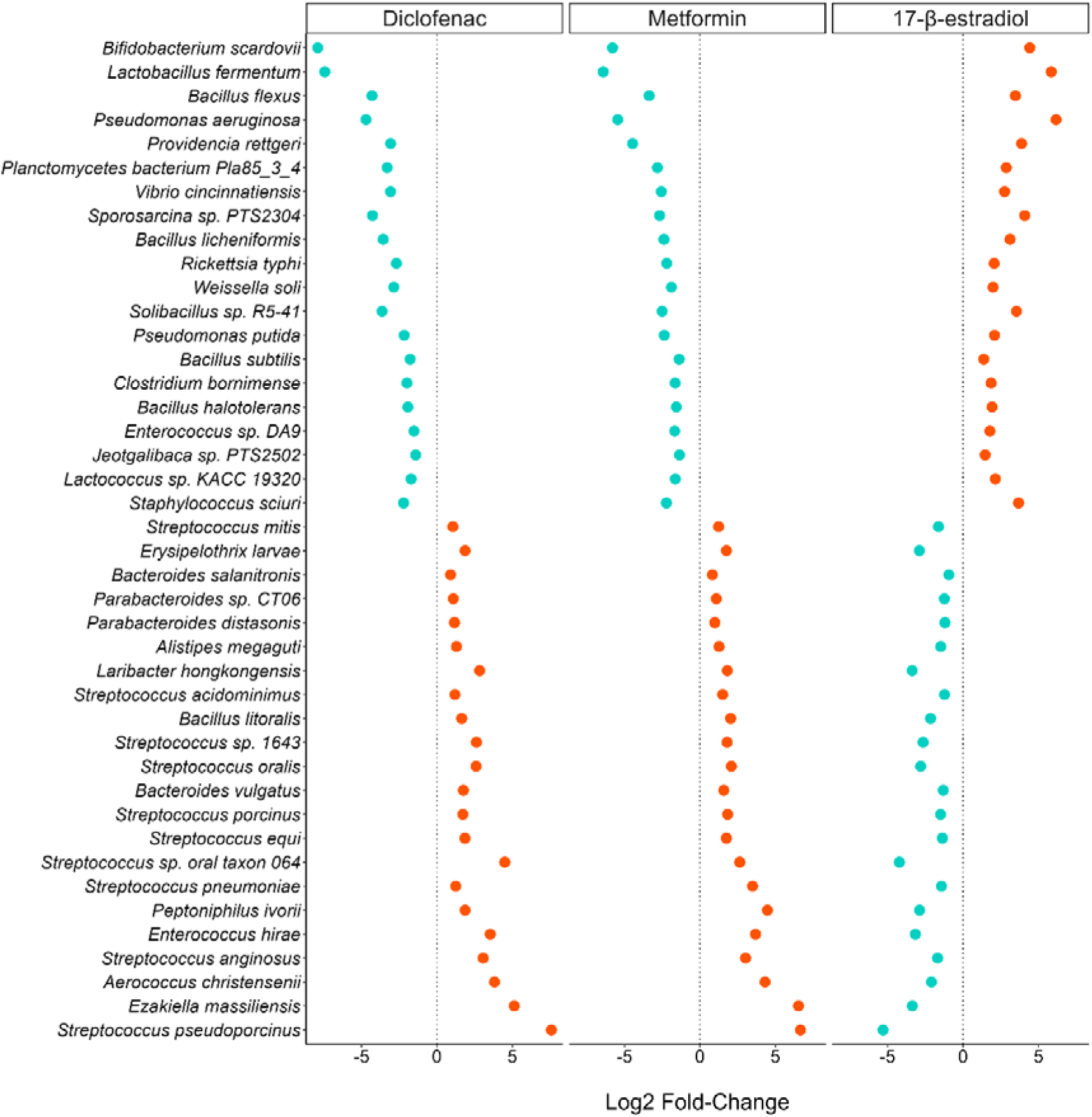
Log2-fold change of species for mixtures compared to ciprofloxacin alone treatment. Each panel consists of pooled data from all ciprofloxacin concentrations across each treatment (i.e. all ciprofloxacin concentrations within the mixture treatments versus all ciprofloxacin concentrations with no NAD, including 0ug/L ciprofloxacin). Those species with a positive change are coloured orange, those with a negative are coloured blue.

## Discussion

Here, we showed that the addition of a NAD at a single spiked concentration can significantly altered the community taxonomy and resistome in a mixed bacterial community, including selecting more strongly for various antibiotic resistance genes. This is despite previous work [45] indicating that these NADs have little to no selective effect when on their own.. We base this on our observations that both growth-based and qPCR-based effect concentrations reduced when NADs were present in mixture with ciprofloxacin. In addition, across all the mixtures, the productivity of the community decreased, indicating that these mixtures are significantly impairing community growth capacity.

Sequencing indicated that mixtures of NADs and ciprofloxacin can select for AMR genes, and can lead to changes to the community composition, although these changes are NAD- and mixture-specific. The concentrations used in this study are sub-MIC, and are much lower than would be used clinically, so the effects we observed might be more profound at higher concentrations of either the antibiotic, NAD, or both.

Firstly, we identified that the richness of the metformin and 17-β-estradiol mixtures showed a decrease in AMR gene richness compared to the ciprofloxacin alone treatment. We suggest that this is due to selection acting upon the AMR genes within these communities, which would lead to increased abundance of less AMR genes.

This is supported by data identifying that these mixtures did show selection for various AMR genes and species. Two of the AMR genes that were significantly altered by the mixtures encode for part of a membrane transport system (*fecE:* an ABC iron transporter [69], and *tolC:* a multi-drug efflux channel [70]). These efflux pump mechanisms may be important in resistance in the metformin and diclofenac mixtures, but not in the 17-β-estradiol mixture. Most intriguing is the finding that the log2-fold changes in species seen differ between diclofenac/metformin mixtures, and the 17-β-estradiol mixture. Coupled with the findings above relating to *fecE* and *tolC,* this might indicate that the 17-β-estradiol mixture is selecting for taxa that are intrinsically resistant or contain unannotated resistance mechanisms. It might also indicate that the diclofenac and metformin mixture is interacting with the community at a molecular level in a similar way. Additionally, we can suggest that the mixtures are acting differently to ciprofloxacin alone with regards to selection for or against different species, with the Gram negative Caulobacterales selected for across the mixtures and negatively selected against by ciprofloxacin alone. This order may be more adaptable towards growth in antibiotic-NAD mixtures, although there is yet no clear evidence as to why. As mentioned, previous work indicated that diclofenac can reduce the MIC of ciprofloxacin [36], and the data presented here indicates that it can also reduce the MSC of ciprofloxacin. The proposed mechanism of diclofenac is inhibition of DNA synthesis [71]. As ciprofloxacin also interferes with DNA synthesis [72, 73], it is likely the combined effects of the two compounds in mixture will significantly reduce replication rate, reflected in the reduction in growth observed in this study. Diclofenac at 10µg/L has also been demonstrated to increase mutation frequency of *E. coli* [19], and to upregulate *sigB*, a gene involved in the stress response [74]. An increased stress response can, in turn, increase mutation rates and integron activation [75, 76]. This might explain some of the *intI1* abundance increases observed in the mixture, and the increases in *tolC* and *fecE* abundances.

Metformin has previously been shown to reduce the MIC of various antibiotics (including ampicillin, doxycycline, and levofloxacin) [37, 77], and it has been suggested that this could be due to the ability of metformin to disrupt cell membranes [37]. This proposed mechanism may allow for increased ciprofloxacin influx into cells, which may lead to selection for any resistance genes present. This might explain why the metformin mixture had the largest increase in *fecE, qnrB*, and *tolC* abundance, since increased ciprofloxacin influx would exert a stronger selection pressure. Metformin acted synergistically with ciprofloxacin, reducing the selective concentration to 10µg/L compared to 40µg/L with ciprofloxacin alone (and it was not selective when tested alone [45]). However, there was an antagonistic effect at 40µg/L. This antagonistic effect may be due to chemical interactions between the two pharmaceuticals. This has been demonstrated to occur previously with ciprofloxacin and zinc [78]. Alternatively, the resistance gene associated with the class I integrons that is under positive selection at 10µg/L may be less beneficial to that community at higher ciprofloxacin concentrations, and a different AMR mechanism is potentially under selection at that point (and unassociated with *intI1*). Future work could aim to understand if this effect holds true for a larger range of concentrations, and whether these pharmaceuticals were indeed interacting chemically.

There is no previous experimental evidence investigating the mixture effect of 17-β-estradiol with an antibiotic. Therefore, these data are presented as novel. Results suggest that 17-β-estradiol may act additively or synergistically with ciprofloxacin to reduce bacterial growth and increase selection for AMR by reducing the minimal selective concentration. In previous work, 17-β-estradiol has been shown to select for *intI1* (from 7µg/L to 5400µg/L), and also selected for metal resistance genes [45]. In this study, the communities exposed to the 17-β-estradiol mixture often responded in the opposite way to the diclofenac and metformin mixtures (for example, the communities exposed to the 17-β-estradiol mixtures showed a decreased *tolC* and *fecE* abundance). Perhaps these communities are relying upon a shift to more tolerant species (as indicated by the fold change in various species), or increased gene expression. Future work could aim to unpick changes to gene expression in response to various NAD and antibiotic mixtures, which might illuminate the changes identified here.

Overall, these results provide some concern for human health. These NADs might be present in the human gut alongside antibiotics, particularly if patients with long-term health conditions (such as diabetes, or those requiring hormone replacement therapy) acquired an infection that required antibiotic treatment. The human gut microbiome can take weeks to recover after antibiotic treatment, and some species never recolonise [79]. In addition to this, after antibiotic treatment there is evidence to suggest that there is an increase in AMR gene abundance in the human gut [80]. Therefore, mixtures of pharmaceuticals in the gut may increase the selective potential of antibiotics, potentially increasing selection for AMR within the gut microbiome. Additionally, of concern is that in the human gut, lower concentrations of ciprofloxacin are required to induce the expression of class I integron [81] (which can contain AMR gene cassettes), and mixtures of pharmaceuticals may exacerbate this. In general, these data suggest that mixtures of NADs and antibiotics lead to selection for AMR at lower concentrations than seen with antibiotics alone, and selectively lead to changes in abundance of species, or specific resistance genes. This could lead to increased selection, maintenance, and dissemination of antibiotic resistance genes between species in the human gut, and increased shedding of these genes and resistant bacteria into the environment.

Furthermore, these pharmaceuticals, alongside others, will be present in the freshwater aquatic, or wastewater environments as micropollutants [23, 31], where they may be affecting growth of the natural communities, and impacting ecosystem functioning. This includes in wastewater treatment plants where they may directly impact wastewater treatment plant functioning if these pharmaceuticals are affecting microorganisms involved in sludge digestion. In our study we found a short-term overall reduction in community growth, but a longer-term increase in Nitrososphaerales abundance, archaea that may play a role in the nitrogen and carbon cycles. Further investigation is needed to determine whether the growth effects are short- or long-term, and what this might mean for ecosystems. These mixtures may lead to increased selection, maintenance, and dissemination of antibiotic resistance genes throughout environmental compartments, or increased input of resistant bacteria to the receiving waters. This then may lead to increased risk of infection or colonisation with resistant strains to people and animals interacting with these environments (e.g. surfers and swimmers in the sea [82]).

Taken together, these data confirm findings that mixtures of NADs and antibiotics can be more selective than the antibiotic or NAD alone. This is of particular concern since previous work [45] has indicated that two of these NADs did not select for AMR in a similar community, and so may have been discounted as potentially selective agents. Many compounds may be disregarded for further study if individually they do not select for AMR. However, data here indicates that inclusion of these compounds in mixture investigations is imperative to understand selection for AMR in more complex mixtures and environments.

## Conclusion

Mixtures of diclofenac, metformin, or 17-β-estradiol with ciprofloxacin both increased the growth inhibitory effects and reduced the selective concentration of ciprofloxacin in a complex bacterial community. Additionally, the mixtures led to selective increases or decreases in specific AMR genes or specific species, some of which are known human pathogens. The effects of antibiotics are traditionally considered in isolation, particularly in terms of selection for AMR. However, antibiotics are present alongside other pharmaceuticals, both in the gut, and in the environment. Further, the data here pertain to simple mixtures, and the effects of more complex mixtures should be considered in future studies, including co-occurring pharmaceuticals which may not be selective in isolation.

## Data Availability

The metagenome data generated and analysed in this study are available at ENA Accession code PRJEB88784. The growth and qPCR data generated and analysed in this study are available at Zenodo DOI: https://doi.org/10.5281/zenodo.15324016. The code used to analyse all the data in the study will be made available upon paper acceptance.

## Authors’ contributions

AH: conceptualisation, data curation, formal analysis, investigation, methodology, visualisation, writing – original draft, writing – review and editing. LZ, EF, JS, BKH: supervision, writing – review and editing. WHG: conceptualisation, methodology, writing – review and editing, supervision. AKM: conceptualisation, funding acquisition, methodology, supervision, writing – review and editing, project administration.

All authors read and approved the final version.

## Acknowledgements

We thank the Exeter Sequencing Service for their work sequencing the Illumina metagenomic sequencing. This project utilised equipment funded by the Wellcome Trust Institutional Strategic Support Fund (WT097835MF), Wellcome Trust Multi User Equipment Award (WT101650MA) and BBSRC LOLA award (BB/K003240/1). The authors would like to acknowledge the use of the University of Exeter’s Advanced Research Computing facilities in carrying out this work.

For the purpose of open access, the authors have applied a Creative Commons Attribution (CC BY) licence to any Author Accepted Manuscript version arising from this submission.

## References

1. Murray CJL, Ikuta KS, Sharara F et al. Global burden of bacterial antimicrobial resistance in 2019: A systematic analysis. The Lancet 2022;399:629–55. 10.1016/S0140-6736(21)02724-0

2. Gullberg E, Cao S, Berg OG et al. Selection of resistant bacteria at very low antibiotic concentrations. PLOS Pathogens 2011;7:e1002158. 10.1371/journal.ppat.1002158

3. Westhoff S, van Leeuwe TM, Qachach O et al. The evolution of no-cost resistance at sub-mic concentrations of streptomycin in streptomyces coelicolor. The ISME Journal 2017;11:1168–78. 10.1038/ismej.2016.194

4. Kraupner N, Ebmeyer S, Bengtsson-Palme J et al. Selective concentration for ciprofloxacin resistance in escherichia coli grown in complex aquatic bacterial biofilms. Environment International 2018;116:255–68. 10.1016/j.envint.2018.04.029

5. Kraupner N, Ebmeyer S, Hutinel M et al. Selective concentrations for trimethoprim resistance in aquatic environments. Environment International 2020;144:106083. 10.1016/j.envint.2020.106083

6. Lundström SV, Ostman M, Bengtsson-Palme J et al. Minimal selective concentrations of tetracycline in complex aquatic bacterial biofilms. Science of The Total Environment 2016;553:587–95. 10.1016/j.scitotenv.2016.02.103

7. Murray AK, Zhang L, Yin X et al. Novel insights into selection for antibiotic resistance in complex microbial communities. mBio 2018;9:e00969–18. 10.1128/mBio.00969-18

8. Sanchez-Cid C, Guironnet A, Keuschnig C et al. Gentamicin at sub-inhibitory concentrations selects for antibiotic resistance in the environment. ISME Communications 2022;2:29. 10.1038/s43705-022-00101-y

9. Stanton IC, Murray AK, Zhang L et al. Evolution of antibiotic resistance at low antibiotic concentrations including selection below the minimal selective concentration. Communications Biology 2020;3:467. 10.1038/s42003-020-01176-w

10. Murray LM, Hayes A, Snape J et al. Co-selection for antibiotic resistance by environmental contaminants. npj Antimicrobials and Resistance 2024;2:9. 10.1038/s44259-024-00026-7

11. Baker-Austin C, Wright MS, Stepanauskas R et al. Co-selection of antibiotic and metal resistance. Trends in Microbiology 2006;14:176–82. 10.1016/j.tim.2006.02.006

12. Maier L, Pruteanu M, Kuhn M et al. Extensive impact of non-antibiotic drugs on human gut bacteria. Nature 2018;555:623–28. 10.1038/nature25979

13. Younis W, Thangamani S, Seleem MN. Repurposing non-antimicrobial drugs and clinical molecules to treat bacterial infections. Curr Pharm Des 2015;21:4106–11. 10.2174/1381612821666150506154434

14. Lu J, Ding P, Wang Y et al. Antidepressants promote the spread of extracellular antibiotic resistance genes via transformation. ISME Communications 2022;2:63. 10.1038/s43705-022-00147-y

15. Wang Y, Lu J, Engelstädter J et al. Non-antibiotic pharmaceuticals enhance the transmission of exogenous antibiotic resistance genes through bacterial transformation. The ISME Journal 2020:2179–96. 10.1038/s41396-020-0679-2

16. Ding P, Lu J, Wang Y et al. Antidepressants promote the spread of antibiotic resistance via horizontally conjugative gene transfer. Environ Microbiol 2022;24:5261–76. 10.1111/1462-2920.16165

17. Wang Y, Lu J, Zhang S et al. Non-antibiotic pharmaceuticals promote the transmission of multidrug resistance plasmids through intra- and intergenera conjugation. The ISME Journal 2021;15:2493–508. 10.1038/s41396-021-00945-7

18. Jin M, Lu J, Chen Z et al. Antidepressant fluoxetine induces multiple antibiotics resistance in escherichia coli via ros-mediated mutagenesis. Environment International 2018;120:421–30. 10.1016/j.envint.2018.07.046

19. Li X, Xue X, Jia J et al. Nonsteroidal anti-inflammatory drug diclofenac accelerates the emergence of antibiotic resistance via mutagenesis. Environmental Pollution 2023;326:121457. 10.1016/j.envpol.2023.121457

20. Wei Z, Wei Y, Li H et al. Emerging pollutant metformin in water promotes the development of multiple-antibiotic resistance in escherichia coli via chromosome mutagenesis. Journal of Hazardous Materials 2022;430:128474. 10.1016/j.jhazmat.2022.128474

21. Hall RJ, Snaith AE, Element SJ et al. Non-antibiotic pharmaceuticals are toxic against escherichia coli with no evolution of cross-resistance to antibiotics. npj Antimicrobials and Resistance 2024;2:11. 10.1038/s44259-024-00028-5

22. Hayes A, Zhang L, Feil E et al. Antimicrobial effects, and selection for amr by non-antibiotic drugs in a wastewater bacterial community. Environment International 2025;199:109490. 10.1016/j.envint.2025.109490

23. aus der Beek T, Weber FA, Bergmann A et al. Pharmaceuticals in the environment--global occurrences and perspectives. Environmental Toxicology and Chemistry 2016;35:823–35. 10.1002/etc.3339

24. Blair BD, Crago JP, Hedman CJ et al. Pharmaceuticals and personal care products found in the great lakes above concentrations of environmental concern. Chemosphere 2013;93:2116–23. 10.1016/j.chemosphere.2013.07.057

25. den Heijer M, Bakker A, Gooren L. Long term hormonal treatment for transgender people. BMJ 2017;359:j5027. 10.1136/bmj.j5027

26. Nichols KC, Schenkel L, Benson H. 17 beta-estradiol for postmenopausal estrogen replacement therapy. Obstetrical & Gynecological Survey 1984;39:230–45. 10.1097/00006254-198404000-00022

27. Ebele AJ, Abou-Elwafa Abdallah M, Harrad S. Pharmaceuticals and personal care products (ppcps) in the freshwater aquatic environment. Emerging Contaminants 2017;3:1–16. 10.1016/j.emcon.2016.12.004

28. Fent K, Weston AA, Caminada D. Ecotoxicology of human pharmaceuticals. Aquatic Toxicology 2006;76:122–59. 10.1016/j.aquatox.2005.09.009

29. UKWIR. Chemicals investigation programme phase 2 database.

30. Stephanie Graumnitz DJ. The database “pharmaceuticals in the environment” update for the period 2017-2020. Report. Umweltbundesamt, 2021

31. Wilkinson JL, Boxall ABA, Kolpin DW et al. Pharmaceutical pollution of the world’s rivers. Proceedings of the National Academy of Sciences 2022;119:e2113947119. 10.1073/pnas.2113947119

32. Hayes A, May Murray L, Catherine Stanton I et al. Predicting selection for antimicrobial resistance in uk wastewater and aquatic environments: Ciprofloxacin poses a significant risk. Environment International 2022;169:107488. 10.1016/j.envint.2022.107488

33. Gil D, Daffinee K, Friedman R et al. Synergistic antibacterial effects of analgesics and antibiotics against staphylococcus aureus. Diagnostic Microbiology and Infectious Disease 2020;96:114967. 10.1016/j.diagmicrobio.2019.114967

34. Chan EWL, Yee ZY, Raja I et al. Synergistic effect of non-steroidal anti-inflammatory drugs (nsaids) on antibacterial activity of cefuroxime and chloramphenicol against methicillin-resistant staphylococcus aureus. Journal of Global Antimicrobial Resistance 2017;10:70–74. 10.1016/j.jgar.2017.03.012

35. Annadurai S, Guha-Thakurta A, Sa B et al. Experimental studies on synergism between aminoglycosides and the antimicrobial antiinflammatory agent diclofenac sodium. Journal of Chemotherapy 2002;14:47–53. 10.1179/joc.2002.14.1.47

36. Hegazy WAH. Diclofenac inhibits virulence of proteus mirabilis isolated from diabetic foot ulcer. African Journal of Microbiology Research 2016;10:733–43.

37. Liu Y, Jia Y, Yang K et al. Metformin restores tetracyclines susceptibility against multidrug resistant bacteria. Advanced Science 2020;7:1902227–27. 10.1002/advs.201902227

38. Tang P-C, Sánchez-Hevia DL, Westhoff S et al. Within-species variability of antibiotic interactions in gram-negative bacteria. mBio 2024;15:e00196–24. doi:10.1128/mbio.00196-24

39. Smith TP, Clegg T, Ransome E et al. High-throughput characterization of bacterial responses to complex mixtures of chemical pollutants. Nature Microbiology 2024;9:938–48. 10.1038/s41564-024-01626-9

40. Klümper U, Recker M, Zhang L et al. Selection for antimicrobial resistance is reduced when embedded in a natural microbial community. The ISME Journal 2019;13:2927–37. 10.1038/s41396-019-0483-z

41. Singh N, Yeh PJ. Suppressive drug combinations and their potential to combat antibiotic resistance. J Antibiot (Tokyo) 2017;70:1033–42. 10.1038/ja.2017.102

42. Tekin E, White C, Kang TM et al. Prevalence and patterns of higher-order drug interactions in escherichia coli. NPJ Syst Biol Appl 2018;4:31. 10.1038/s41540-018-0069-9

43. Sutradhar I, Ching C, Desai D et al. Effects of antibiotic interaction on antimicrobial resistance development in wastewater. Sci Rep 2023;13:7801. 10.1038/s41598-023-34935-w

44. Gillings MR, Gaze WH, Pruden A et al. Using the class 1 integron-integrase gene as a proxy for anthropogenic pollution. The ISME Journal 2015;9:1269–79. 10.1038/ismej.2014.226

45. Hayes A, Zhang L, Feil E et al. Antimicrobial effects, and selection for amr by non-antibiotic drugs on bacterial communities. bioRxiv 2024:2024.04.23.590690. 10.1101/2024.04.23.590690

46. Ewels P, Magnusson M, Lundin S et al. Multiqc: Summarize analysis results for multiple tools and samples in a single report. Bioinformatics 2016;32:3047–48. 10.1093/bioinformatics/btw354

47. Bonin N, Doster E, Worley H et al. Megares and amr++, v3.0: An updated comprehensive database of antimicrobial resistance determinants and an improved software pipeline for classification using high-throughput sequencing. Nucleic Acids Res 2023;51:D744–d52. 10.1093/nar/gkac1047

48. Pal C, Bengtsson-Palme J, Rensing C et al. Bacmet: Antibacterial biocide and metal resistance genes database. Nucleic Acids Res 2014;42:D737–D43. 10.1093/nar/gkt1252

49. Florensa AF, Kaas RS, Clausen P et al. Resfinder - an open online resource for identification of antimicrobial resistance genes in next-generation sequencing data and prediction of phenotypes from genotypes. Microb Genom 2022;8 10.1099/mgen.0.000748

50. Alcock BP, Huynh W, Chalil R et al. Card 2023: Expanded curation, support for machine learning, and resistome prediction at the comprehensive antibiotic resistance database. Nucleic Acids Res 2023;51:D690–d99. 10.1093/nar/gkac920

51. Wood DE, Lu J, Langmead B. Improved metagenomic analysis with kraken 2. Genome Biology 2019;20:257. 10.1186/s13059-019-1891-0

52. O’Leary NA, Wright MW, Brister JR et al. Reference sequence (refseq) database at ncbi: Current status, taxonomic expansion, and functional annotation. Nucleic Acids Res 2016;44:D733–45. 10.1093/nar/gkv1189

53. Pruitt KD, Tatusova T, Maglott DR. Ncbi reference sequences (refseq): A curated non-redundant sequence database of genomes, transcripts and proteins. Nucleic Acids Res 2007;35:D61–5. 10.1093/nar/gkl842

54. McMurdie PJ, Holmes S. Phyloseq: An r package for reproducible interactive analysis and graphics of microbiome census data. PLOS ONE 2013;8:e61217. 10.1371/journal.pone.0061217

55. Author. Metagenomeseq: Statistical analysis for sparse high-throughput sequncing [Computer software]. http://www.cbcb.umd.edu/software/metagenomeSeq. 2031.

56. Paulson JN, Stine OC, Bravo HC et al. Differential abundance analysis for microbial marker-gene surveys. Nature Methods 2013;10:1200–02. 10.1038/nmeth.2658

57. Author. R: A language and environment for statistical computing [Computer software]. R Foundation for Statistical Computing. 2019.

58. Wickham H. Ggplot2: Elegant graphics for data analysis. Springer-Verlag New York, 2016.

59. Author. Metbrewer: Color palettes inspired by works at the metropolitan museum of art [Computer software]. 2022.

60. Hartig F. Dharma: Residual diagnostics for hierarchical (multi-level/mixed) regression models. R package version 03 2020;3

61. Murray AK, Stanton IC, Wright J et al. The selection end points in communities of bacteria (select) method: A novel experimental assay to facilitate risk assessment of selection for antimicrobial resistance in the environment. Environmental Health Perspectives 2020;128:107007. doi:10.1289/EHP6635

62. Sprouffske K, Wagner A. Growthcurver: An r package for obtaining interpretable metrics from microbial growth curves. BMC Bioinformatics 2016;17:172. 10.1186/s12859-016-1016-7

63. Bates D, Machler M, Bolker B et al. Fitting linear mixed-effects models using {lme4}. Journal of Statistical Software 2015:1–48. 10.18637/jss.v067.i01

64. Lenth R, Singmann H, Love J et al. Emmeans: Estimated marginal means, aka least-squares means. R package version 1 (2018).

65. Shannon CE. A mathematical theory of communication. The Bell system technical journal 1948;27:379–423.

66. Dixon P. Vegan, a package of r functions for community ecology. Journal of Vegetation Science 2003;14:927–30. 10.1111/j.1654-1103.2003.tb02228.x

67. Love MI, Huber W, Anders S. Moderated estimation of fold change and dispersion for rna-seq data with deseq2. Genome Biology 2014;15:550. 10.1186/s13059-014-0550-8

68. Greenfield BK, Shaked S, Marrs CF et al. Modeling the emergence of antibiotic resistance in the environment: An analytical solution for the minimum selection concentration. Antimicrob Agents Chemother 2018;62 10.1128/aac.01686-17

69. Van Hove B, Staudenmaier H, Braun V. Novel two-component transmembrane transcription control: Regulation of iron dicitrate transport in escherichia coli k-12. J Bacteriol 1990;172:6749–58. 10.1128/jb.172.12.6749-6758.1990

70. Paulsen IT, Park JH, Choi PS et al. A family of gram-negative bacterial outer membrane factors that function in the export of proteins, carbohydrates, drugs and heavy metals from gram-negative bacteria. FEMS Microbiology Letters 1997;156:1–8. 10.1111/j.1574-6968.1997.tb12697.x

71. Dastidar SG, Ganguly K, Chaudhuri K et al. The anti-bacterial action of diclofenac shown by inhibition of DNA synthesis. International Journal of Antimicrobial Agents 2000;14:249–51. 10.1016/S0924-8579(99)00159-4

72. Nakamura S, Nakamura M, Kojima T et al. Gyra and gyrb mutations in quinolone-resistant strains of escherichia coli. Antimicrob Agents Chemother 1989;33:254–5. 10.1128/aac.33.2.254

73. Yoshida H, Kojima T, Yamagishi J et al. Quinolone-resistant mutations of the gyra gene of escherichia coli. Molecular Genetics and Metabolism 1988;211:1–7. 10.1007/bf00338386

74. Riordan JT, Dupre JM, Cantore-Matyi SA et al. Alterations in the transcriptome and antibiotic susceptibility of staphylococcus aureus grown in the presence of diclofenac. Ann Clin Microbiol Antimicrob 2011;10:30–30. 10.1186/1476-0711-10-30

75. Guerin É, Cambray G, Sanchez-Alberola N et al. The sos response controls integron recombination. Science 2009;324:1034–34. doi:10.1126/science.1172914

76. Hocquet D, Llanes C, Thouverez M et al. Evidence for induction of integron-based antibiotic resistance by the sos response in a clinical setting. PLOS Pathogens 2012;8:e1002778. 10.1371/journal.ppat.1002778

77. Masadeh MM, Alzoubi KH, Masadeh MM et al. Metformin as a potential adjuvant antimicrobial agent against multidrug resistant bacteria. Clin Pharmacol 2021;13:83–90. 10.2147/CPAA.S297903

78. Vos M, Sibleyras L, Lo LK et al. Zinc can counteract selection for ciprofloxacin resistance. FEMS Microbiology Letters 2020;367:fnaa038. 10.1093/femsle/fnaa038

79. Palleja A, Mikkelsen KH, Forslund SK et al. Recovery of gut microbiota of healthy adults following antibiotic exposure. Nature Microbiology 2018;3:1255–65. 10.1038/s41564-018-0257-9

80. MacPherson CW, Mathieu O, Tremblay J et al. Gut bacterial microbiota and its resistome rapidly recover to basal state levels after short-term amoxicillin-clavulanic acid treatment in healthy adults. Scientific Reports 2018;8:11192. 10.1038/s41598-018-29229-5

81. Baltazar M, Bourgeois-Nicolaos N, Larroudé M et al. Activation of class 1 integron integrase is promoted in the intestinal environment. PLOS Genetics 2022;18:e1010177. 10.1371/journal.pgen.1010177

82. Leonard AFC, Zhang L, Balfour AJ et al. Exposure to and colonisation by antibiotic-resistant e. Coli in uk coastal water users: Environmental surveillance, exposure assessment, and epidemiological study (beach bum survey). Environment International 2018;114:326–33. 10.1016/j.envint.2017.11.003

