## Supplementary File for "Common non-antibiotic drugs enhance selection for antimicrobial resistance in mixture with ciprofloxacin"

**Section 1. Full details on model outputs for AUC**

For the diclofenac and ciprofloxacin mixture, both diclofenac treatment (p<0.001), and ciprofloxacin concentration (p<0.001) were significant in the model (F_3,92_=39.86, p=<2.2e-16). The AUC of diclofenac at the LOEC was significantly lower than the AUC of the NOEC of diclofenac mixture (p<0.0001) and the AUC of ciprofloxacin alone (p<0.0001). There was no difference in the AUC between the ciprofloxacin alone treatment, and the diclofenac NOEC and ciprofloxacin treatment (p=0.94).

For the metformin mixture, again both metformin treatment (p<0.001) and ciprofloxacin concentration (p<0.001) significantly affected the AUC, and the model was significant (F_3, 92_=50.74, p=< 2.2e-16). The metformin LOEC mixture significantly reduced the AUC in mixture with ciprofloxacin when compared to growth in ciprofloxacin alone (p=<0.0001), and ciprofloxacin and metformin NOEC mixture (p=<0.0001) (Figure 7). The AUC did not differ between growth in ciprofloxacin alone and ciprofloxacin and metformin NOEC (p=0.60).

For the 17-β-estradiol mixture, both 17-β-estradiol treatment (p=0.001), and ciprofloxacin concentration (p=0.011), significantly affected the AUC, and the model was significant (F_3, 92_=31.86, p=3.249e-14). The LOEC of 17-β-estradiol and ciprofloxacin mixture had a significantly reduced AUC when compared to growth in only ciprofloxacin (p<0.0001) and growth in ciprofloxacin and the NOEC of 17-β-estradiol (p<0.0001). The AUC did not differ between growth in ciprofloxacin alone and ciprofloxacin and 17-β-estradiol NOEC (p=0.94).

**Section 2. Full details of model outputs for genes of interest**

***Aph3-DPRIME***

Aph3-DPRIME abundance is reduced by both ciprofloxacin concentration (main effect: F_1, 464_=15.31, p<0.001), and mixture type (main effect: F_3, 464_=7.03, p<0.001), and by the interaction of both ciprofloxacin concentration and mixture type (interaction effect: F_3, 464_=2.69, p=0.045).

***Aph6***

*Aph6* abundance is altered by both mixture type (main effect: F_3,460_ = 16.7, p<0.0001), and the interaction between mixture and the unimodal concentration term (interaction effect: F_3,460_=4.2, p=0.006).

***FecE***

*FecE* abundance altered with both ciprofloxacin concentration (main effect: F_1,110_=20.65, p<0.001), and mixture type (main effect: F_3, 110_=3.25, p=0.025), and by the interaction of both ciprofloxacin concentration and mixture type (interaction effect: F_3, 110_=12.64, p<0.001). Whereby the 17-β-estradiol and ciprofloxacin treatments show a reduced *fecE* abundance, and the diclofenac metformin mixture show a strong increased *fecE* abundance with increasing ciprofloxacin concentration.

***QnrB***

*QnrB* abundance increased with ciprofloxacin concentration (main effect F_1, 51_=156.9, p<0.001), and mixture type (main effect: F_3,51_=2.8, p=0.047). However, the interaction between ciprofloxacin concentration and mixture type was not significant but trended towards being so (interaction effect: F_3,51_=2.7, p=0.052). The trend towards significant with the interaction term is driven by the 17-β-estradiol mixture, which showed the smallest rate of increase in *qnrB* abundance than the other mixture types, despite all mixture types and ciprofloxacin alone treatments showing an increase in *qnrB* abundance with ciprofloxacin concentration.

***tetA***

*TetA* showed a significant change with both the linear concentration effect (main effect: F_1,342_=89, p<0.001) and the unimodal term of concentration (main effect: F_1,342_=57.9, p<0.001). *TetA* also altered with mixture type (main effect: F_3,342_=5.6, p<0.001), and the interaction between mixture and the linear concentration (interaction effect: F_3,342_=4.1, p=0.006), and the interaction between mixture and the unimodal term of concentration (interaction effect: F_3,342_=6.5, p<0.001).

***tetQ***

*TetQ* abundance was not affected by ciprofloxacin concentration, but mixture type significantly altered *tetQ* abundance (F_3, 291_=4.96, p=0.0023).

***toLC***

*TolC* tended to change with concentration (main effect: F_1,169=_10.89, p=0.0012), and was different with mixture treatments (main effect F_3,169_=4.0, p=0.009). The interaction between concentration and mixture was also significant (interaction effect: F_3,169_=11.5, p<0.001), where 17-β-estradiol mixture decreased *tolC* abundance with ciprofloxacin concentration, but ciprofloxacin alone mixture showed a slight increased *tolC* abundance with ciprofloxacin concentration, and the diclofenac and metformin mixtures had the largest increase in the *tolC* abundance.

**Supplementary Figures**

**Supplementary Figure 1.** **QPCR plots with both day 0 and day 7 data. IntI1 prevalence as a function of ciprofloxacin concentration in both ciprofloxacin alone (A) or in combination with diclofenac (B), metformin (C), or 17-β-estradiol (D). Five biological replicates shown. Significant differences of day seven prevalences are shown from pairwise comparisons to the day seven prevalence at 0µg/L. NS = non-significant, * = p<0.05, ** = p<0.01, *** = p<0.001.**

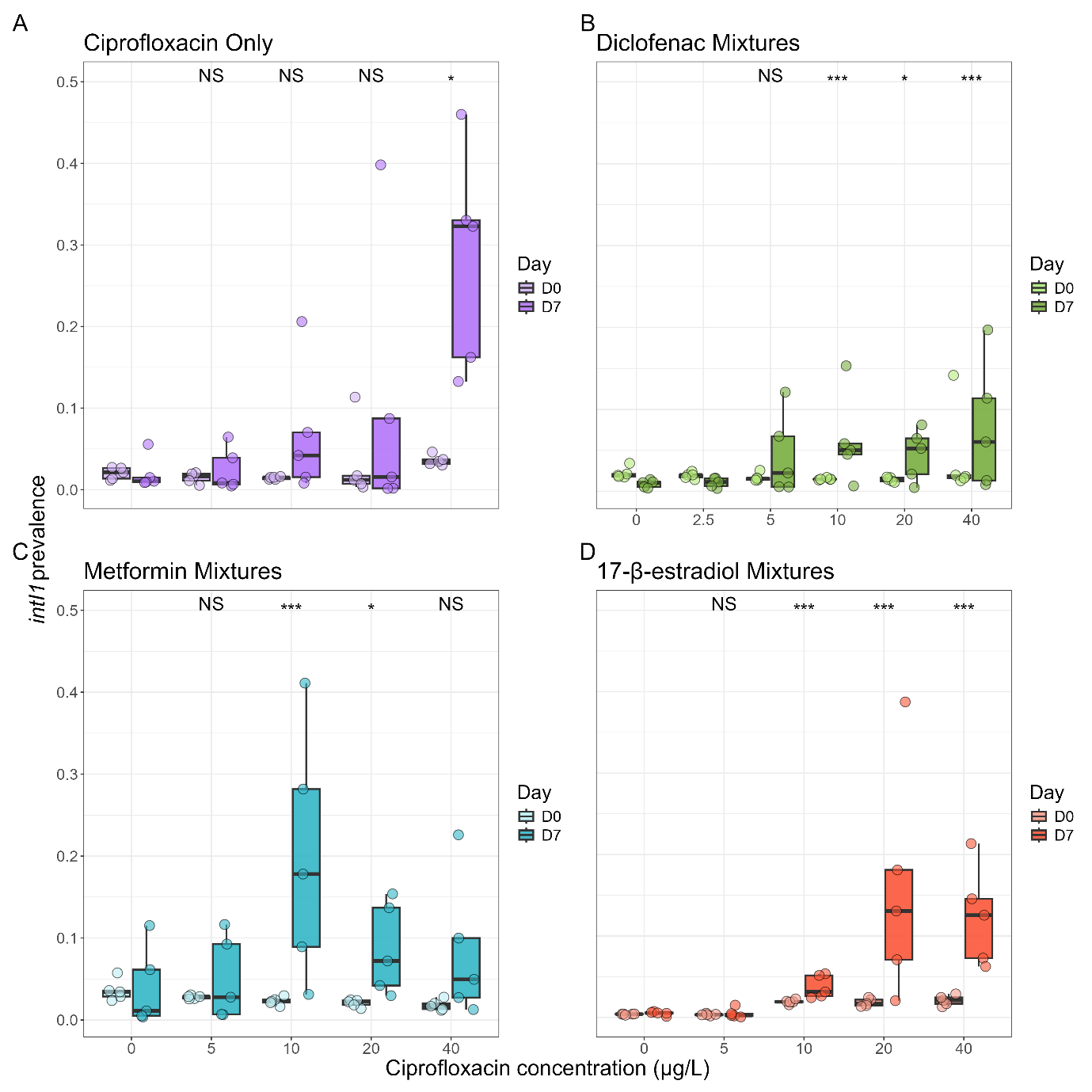

**Supplementary Figure 2. NMDS plot showing Bray-Curtis ordination of resistome of evolved treatments. Dotted ellipses indicate 95% confidence interval for each mixture type.**

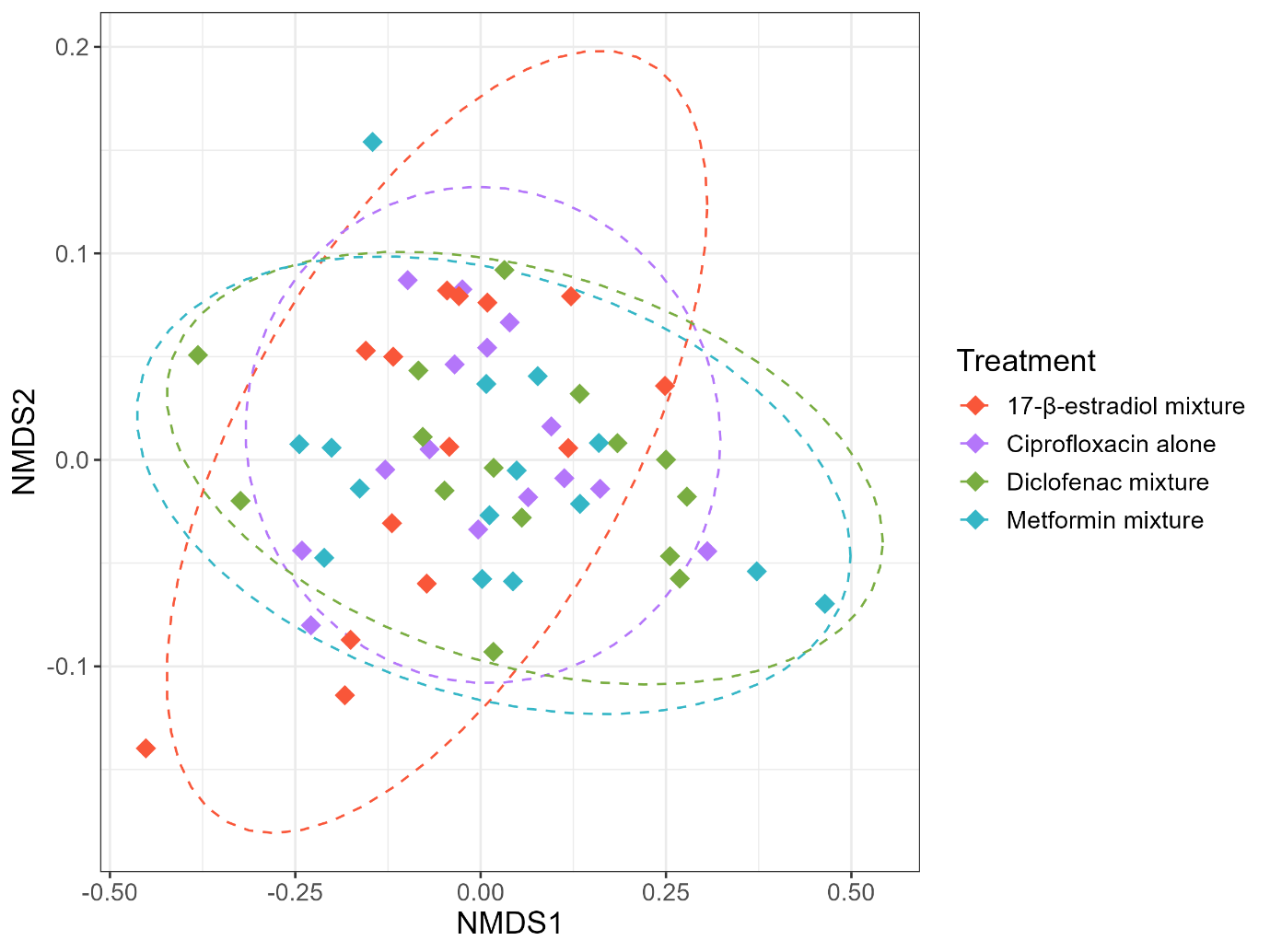

**Supplementary Figure 3. NMDS plot showing Bray-Curtis ordination of taxonomy of evolved treatments. Dotted ellipses indicate 95% confidence interval for each mixture type.**

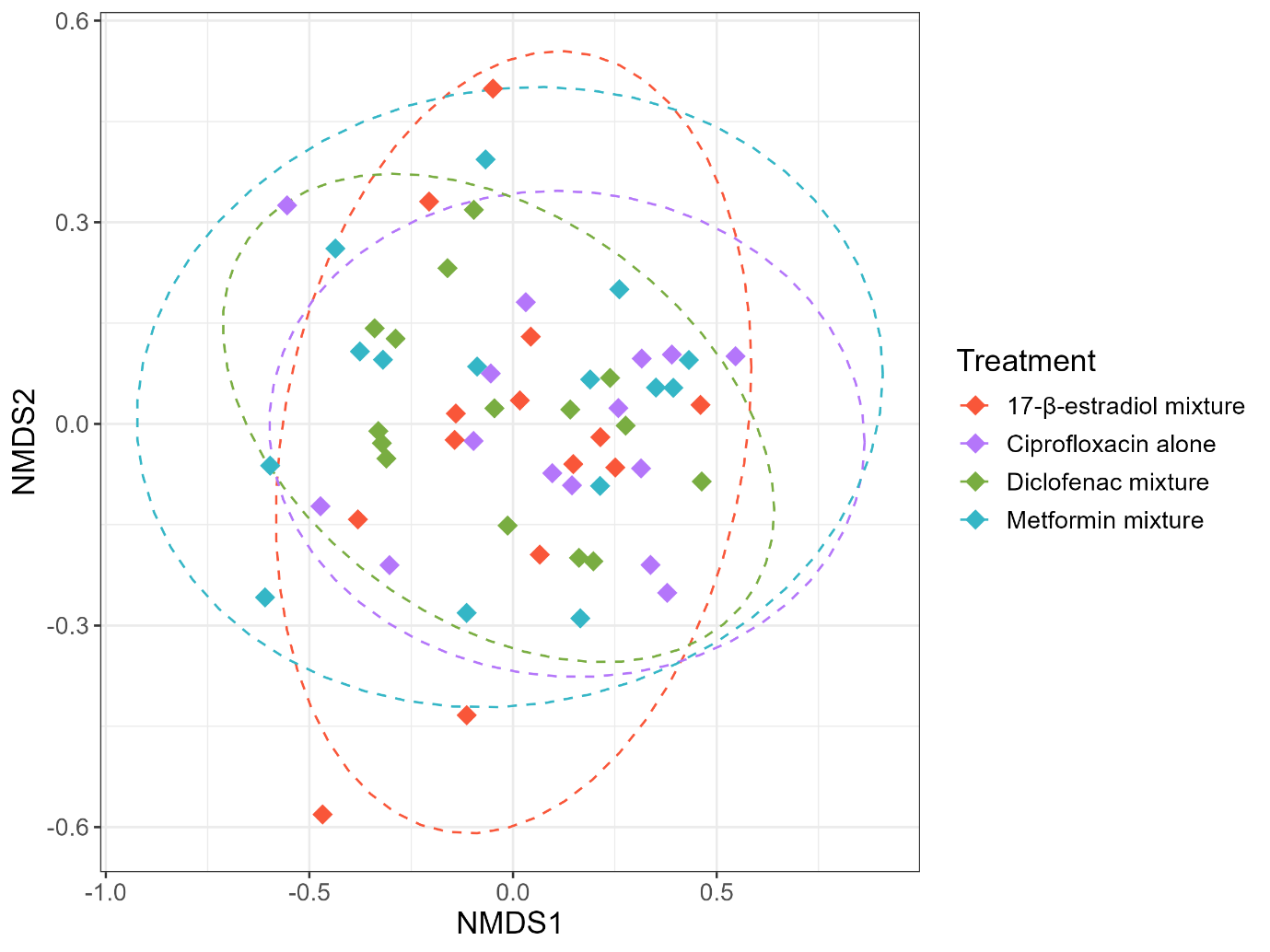

**Supplementary Figure 4. Stacked bar plots showing the relative abundance of A) high abundance genera and B) all phyla.**

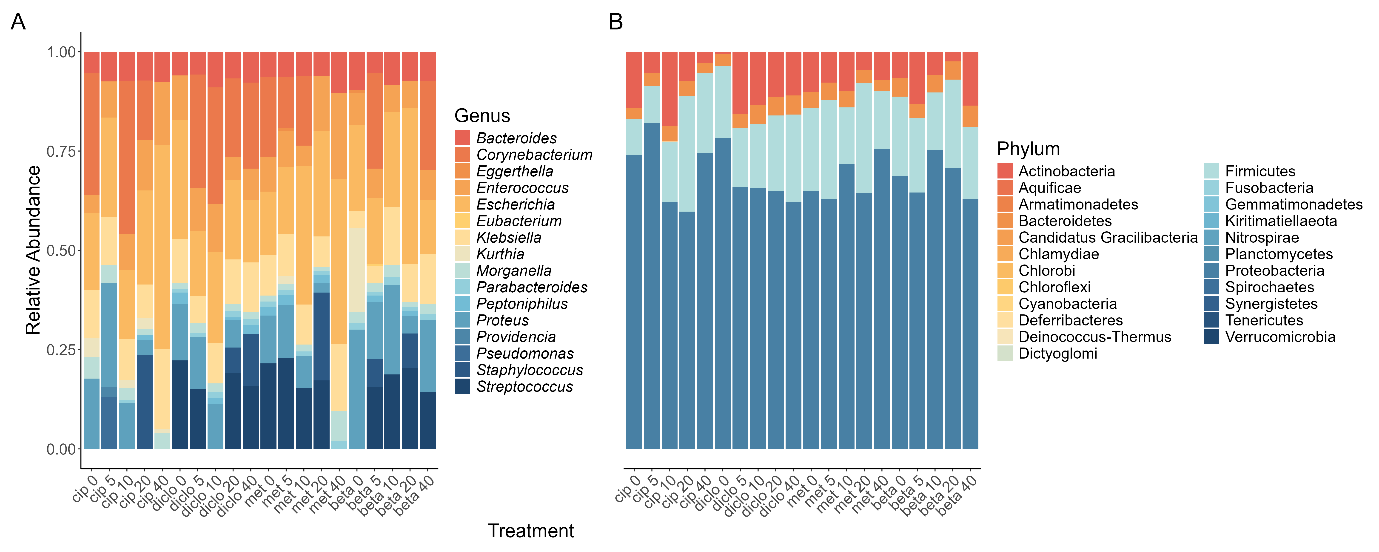

**Supplementary Figure 5. Log2 fold change of species in the A) the diclofenac mixture, B) the metformin mixture, or C) the 17-β-estradiol mixture, compared to the ciprofloxacin alone treatment.**

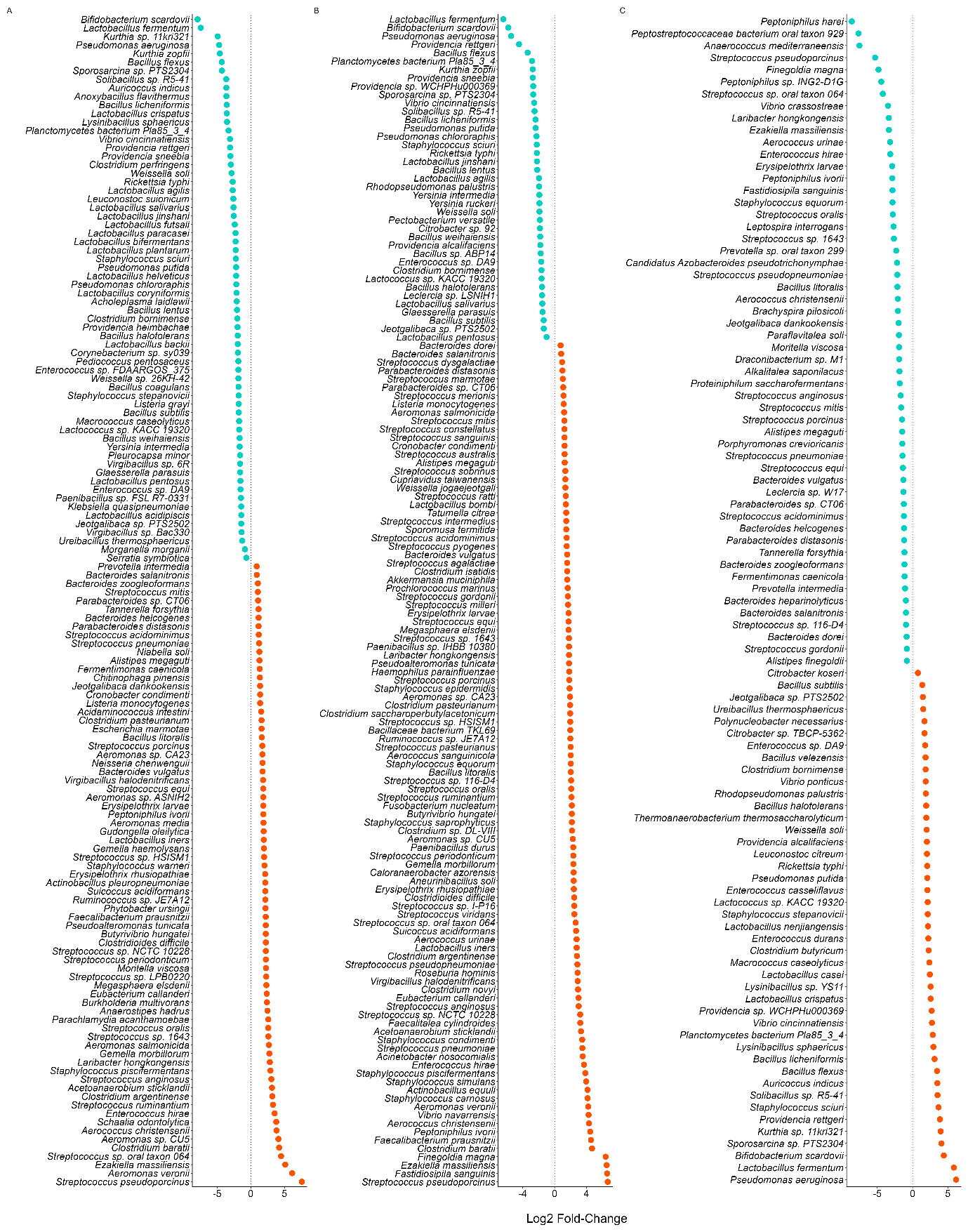

**Supplementary Table 1. Primer and gBlock sequences used in qPCR assays**

| **Target Name** | **Sequence** | | **Reference** |
| --- | --- | --- | --- |
| *16S* rRNA | Primer | CGGTGAATACGTTCYCGG  GGWTACCTTGTTACGACT | Suzuki et al., 2000. 10.1128/aem.66.11.4605-4614.2000 |
|  | gBlock | GCTACGGTGAATACGTTCCCGGGCCTTGTACACACCGCCCGTCACACCATGGGAGTGGGTTGCAAAAGAAGTAGGTAGCTTAACCTTCGGGAGGGCGCTTACCACTTTGTGATTCATGACTGGGGTGAAGTCGTAACAAGGTAACCGTAGG |  |
| *intI1* | Primer | GCCTTGATGTTACCCGAGAG  GATCGGTCGAATGCGTGT | Barraud et al., 2010. 10.1093/jac/dkq167 |
|  | gBlock | catGGCCTTGATGTTACCCGAGAGCTTGGCACCCAGCCTGCGCGAGCAGCTGTCGCGTGCACGGGCATGGTGGCTGAAGGACCAGGCCGAGGGCCGCAGCGGCGTTGCGCTTCCCGACGCCCTTGAGCGGAAGTATCCGCGCGCCGGGCATTCCTGGCCGTGGTTCTGGGTTTTTGCGCAGCACACGCATTCGACCGATCCATA |  |

**Supplementary Table 2. Model outputs of Kruskal-Wallis tests of significance for all fluoroquinolone resistance genes in all samples in this study.**

| **Resistance Gene** | **Adjusted p value** |
| --- | --- |
| *qnrB* | 0.00051 |
| *parEF* | 0.13 |
| *gyrA* | 1.00 |
| *gyrB* | 1.00 |
| *parC* | 1.00 |

**Supplementary Table 3. Resistance genes that were significantly different in at least one treatment, across the three NAD and ciprofloxacin mixtures.**

| **Resistance Gene** | **Adjusted p value** |
| --- | --- |
| *CTX* | <0.0001 |
| *QNRB* | 0.0043 |
| *TOLC* | 0.040 |
| *FECE* | 0.013 |
| *SILC* | 0.0090 |
| *APH3-DPRIME* | <0.0001 |
| *TETA* | <0.0001 |
| *APH6* | <0.0001 |
| *TETQ* | 0.0028 |
| *SULII* | 0.0090 |
